# Further investigation of the potential anti-neoplastic, anti-inflammatory and immunomodulatory actions of phenoxybenzamine using the Broad Institute CLUE platform

**DOI:** 10.1101/767392

**Authors:** Mario A. Inchiosa

## Abstract

Previous clinical studies with the FDA-approved alpha-adrenergic antagonist, phenoxybenzamine, showed apparent efficacy to reverse the symptoms and disabilities of the neuropathic condition, Complex Regional Pain Syndrome; also, the anatomic spread and intensity of this syndrome has a proliferative character and it was proposed that phenoxybenzamine may have an anti-inflammatory, immunomodulatory mode of action. A previous study gave evidence that phenoxybenzamine had anti-proliferative activity in suppression of growth in several human tumor cell cultures. The same report demonstrated that the drug possessed significant histone deacetylase inhibitory activity. Utilizing the Harvard/Massachusetts Institute of Technology Broad Institute genomic database, CLUE, the present study suggests that the gene expression signature of phenoxybenzamine in malignant cell lines is consistent with anti-inflammatory/immunomodulatory activity and suppression of tumor expansion by several possible mechanisms of action. Of particular note, phenoxybenzamine demonstrated signatures that were highly similar to those with glucocorticoid agonist activity. Also, gene expression signatures of phenoxbenzamine were consistent with several agents in each case that were known to suppress tumor proliferation, notably, protein kinase C inhibitors, Heat Shock Protein inhibitors, epidermal growth factor receptor inhibitors, and glycogen synthase kinase inhibitors. Searches in CLUE also confirmed the earlier observations of strong similarities between gene expression signatures of phenoxybenzamine and several histone deacetylase inhibitors.

## 1. Introduction

A previous study explored a possible anti-proliferative capacity of the drug, phenoxybenzamine (PBZ), as a basis for its apparent therapeutic potential in the treatment of chronic neuropathic pain in the Complex Regional Pain Syndrome (CRPS) (1). The BroadBuild02, Broad Institute Harvard/Massachusetts Institute of Technology (MIT) Molecular Signature Database (MSigDB) and the associated Connectivity Map (CMap) software were explored for a possible basis for this effect (2, 3). Gene expression signatures for PBZ on the CMap platform showed appreciable similarity to classical histone deacetylase (HDAC) inhibitors. An extensive comparison of PBZ with suberanilohydroxamic acid (SAHA) and trichostatin A (TSA) demonstrated gene expression enrichment scores (connectivity) for PBZ that compared equal to or greater than those for SAHA and TSA for several gene expression signatures in MSigDB that are associated with histone modifications (1).

For example, this was true for gene expression signatures from ultraviolet exposure of the epidermis in the 100 to 280 nm wavelength range (UVC class radiation); such exposures are associated with trichothiodystrophy syndrome and xeroderma pigmentosum (4). PBZ also showed highly similar gene expression signatures as those for the histone modifying effects of SAHA and TSA on the viral replication process of human cytomegalovirus; these effects on histones have a “therapeutic” value in the application of oncolytic cancer therapy that is herpes-based (5–7). A third area of strong agreement among PBZ, SAHA and TSA related to the similarities among their gene expression signatures to those resulting from ultraviolet radiation of normal epidermal keratinocytes in the 280 to 320 nm range (UVB class radiation). This class of radiation can result in the development of squamous cell or basal cell carcinoma. TSA has been demonstrated to aid in the recovery of markers of keratinocyte differentiation following UVB radiation (8). A fourth area of overlap of gene enrichment scores among HDAC inhibitors and PBZ related to the known inhibitory effects of TSA, SAHA, valproic acid and sodium butyrate on colon cancer cell lines (9).

In vitro assays in the earlier work confirmed HDAC inhibitory activity for PBZ in several classes of HDACs, with the greatest inhibition upon HDACs 5, 6 and 9(1). In the same communication, inhibitory effects of PBZ were demonstrated on the proliferation of several human tumor cell lines; the most sensitivity was observed in non-small-cell lung, lymphoma, myeloma, ovarian, and prostate cell cultures.

The present study concerns further investigation of PBZ gene expression profiles in the new cloud-based version of the Broad Institute gene expression database, the CLUE platform; this database continues to be identified as the Connectivity Map (CMap) (10). However, the new platform is a major expansion of the catalog of cellular gene signature responses of human malignant and premalignant cells to chemical and genetic perturbation.

The analyses in the present study have identified a number of gene expression signatures for PBZ that are closely similar to a large group of glucocorticoids, drugs that are known to have anti-inflammatory and immunomodulatory actions; these agents have a steroid chemical structure, but PBZ has no elements of such a structure (Fig 1). PBZ also had strong gene signature connectivity with several groups of agents that have well-identified roles in the inhibition of signaling pathways for malignant expansion. This appears to be supportive of the previous observations of inhibition of growth in malignant tumor cell cultures (1). The CLUE CMap also confirms the similarities among gene expression signatures for PBZ and HDAC inhibitors in the expanded group of 9 malignant and premalignant cell lines. The present studies of potential anti-proliferative, anti-inflammatory and immunomodulatory activity for PBZ would appear to support the previous observations and help to develop hypotheses for repurposing PBZ for new therapeutic applications. Finally, recent studies by other investigators, which are enumerated herein, are consistent with a broadening perspective about the potential therapeutic attributes of the drug.

**Fig 1.**
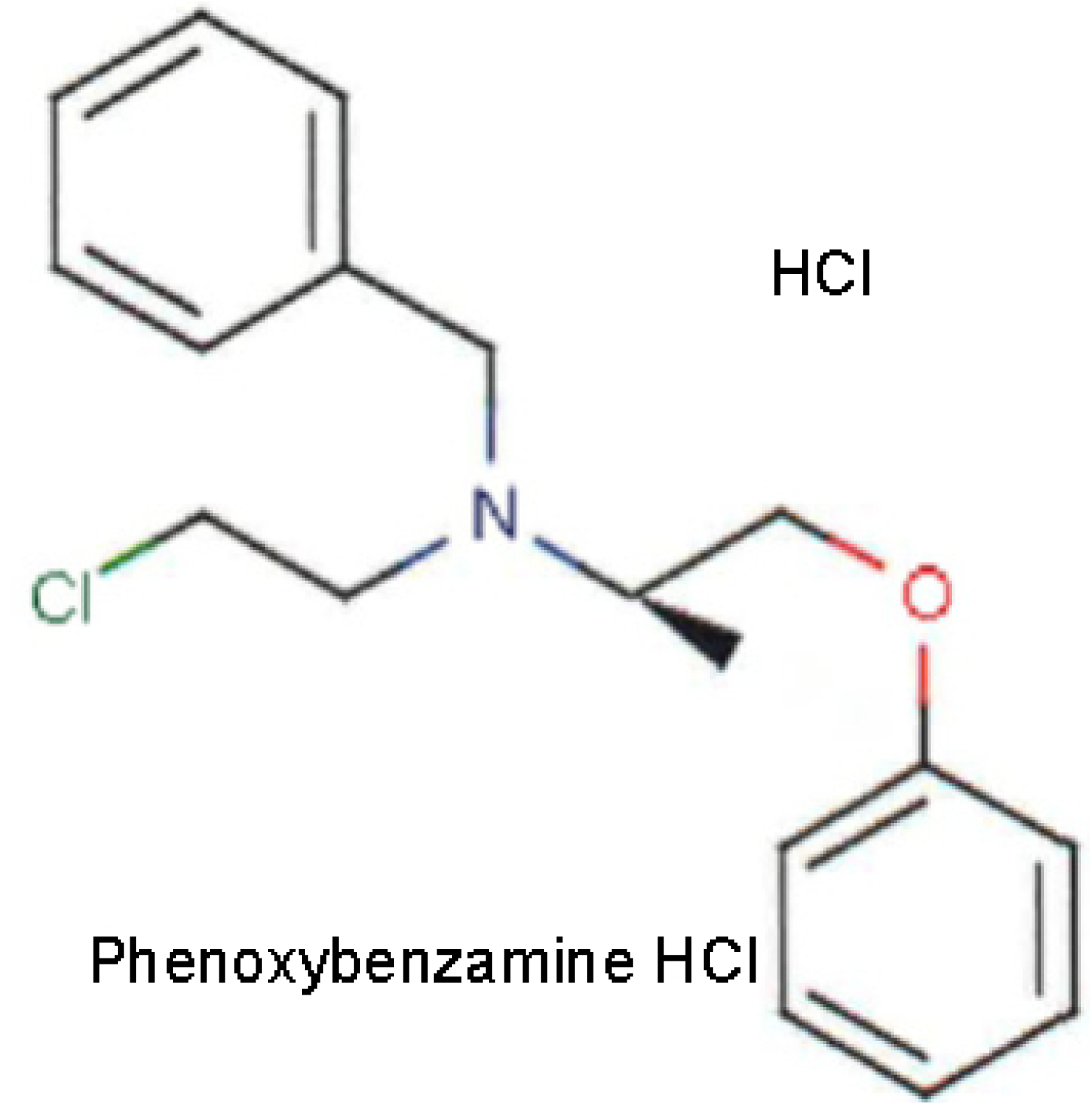
Chemical structure of phenoxybenzamine HCl

## 2. Methods

Both the earlier online Broad Institute CMap genomic dataset and the new cloud-based CLUE platform are based on the gene expression signatures that result from the perturbation of actively proliferating cells that are malignant and, in one case, premalignant. The database and associated software are accessible at https://clue.io. The drugs, chemical agents and genetic inputs are referred to as, “perturbagens” (2, 10). The present analyses focused on gene-expression signatures located in the “TOUCHSTONE” dataset (accessed in the “Tools” menu) that was selected because it represents a set of thoroughly annotated small molecule perturbagens that would be expected to be relevant to comparisons with the possible effect of PBZ; see: https://clue.io/connectopedia/tag/TOUCHSTONE. The Touchstone dataset contains a total of approximately 8,400 perturbagens that have produced gene signatures that were generated from testing on a panel of 9 cell lines; those cell lines are detailed in Table 1. In almost all instances, the perturbagens were tested at a concentration of 10 µM, allowing for the comparison of the strength of the connectivity of gene expression between agents at equimolar concentrations; this included the instances for PBZ, as well.

**Table 1.** Cell Lines Profiled in the TOUCHSTONE Database

| Cell Line | Description |
| --- | --- |
| A375 | Human malignant melanoma |
| A549 | Human non-small cell carcinoma |
| HA1E | Human kidney epithelial immortalized |
| HCC515 | Human non-small cell lung adenocarcinoma |
| HEPG2 | Human hepatocellular carcinoma cell line |
| MCF7 | Human breast adenocarcinoma |
| PC3 | Human prostate adenocarcinoma |
| VCAP | Human metastatic prostate cancer |
| HT29 | Human colorectal adenocarcinoma |

It should be noted that the dataset in the earlier online platform of CMap was based almost entirely on perturbations in only 3 tumor cell lines, including MCF7 (human breast adenocarcinoma), PC3 (human prostate adenocarcinoma) and HL60 (human promyeloblast); the first two cell lines are among the 9 in the current CLUE platform. In fact, in the case of PBZ, it was only screened with the MCF7 and PC3 cell lines.

The scoring value obtained in the gene expression analyses is termed “tau;” it ranges from 100 to −100 and is a measure of the connectivity between the gene expression signature of the perturbagen of interest (PBZ in our searches) and those of the other 8400 perturbagens in the database. A positive tau indicates a relative similarity between two perturbagens or group of perturbagens, while a negative score indicates relative opposing gene signatures. Thus, for example, a tau score of 95 indicates that only 5% of the signatures in the database had connectivities that were higher than those to PBZ; see: https://clue.ie/connectipedia/connectivity_scores.

The Touchstone site returns two data formats in response to a query about the similarities or differences between a perturbagen of interest and the other perturbagens in the database. The compound being searched against all other entries in the Touchstone database is termed the “INDEX.” A “heatmap” format presents the connectivity score between the query perturbagen (PBZ) and each of the reference perturbagens in the database for each of the 9 cells lines that have been studied. The second format is the “detailed list” output. In our experience, this output provided the most convenient access to information that relates to the primary focus of the present studies. The detailed list format also includes the protein targets of the individual perturbagens. Both formats provide comparison of the gene signatures of PBZ with members of the four perturbagen classes in the Touchstone database. Those classes are identified as: Chemical compound/pharmacologic agent (CP); Gene knock-down (KD); Gene over-expression (OE); and perturbagen class (PCL). In the heat map format, PBZ signatures were compared with the entire Touchstone database of perturbagens (approximately 8400). Since our primary interest was to search for drugs and chemicals that had recognized mechanisms of action that might strengthen previous findings and support hypotheses for the repurposing of PBZ therapeutically, we focused our searches in the detailed list format to the CP agents; this represented approximately 2400 compounds.

The heatmap output that is presented gives the calculated *median* tau score (connectivity) for the 9 cell lines and a “summarization” measure. For all other results, the connectivity scores that are presented are the summarization measure. This measure is calculated across the 9 cell lines. It is valuable since it provides a measure of the consistency of perturbations from one cell line to another. The algorithms for the calculation of tau and the summarization measure, and for the associated formulas, are presented at: https://clue.io/connectopedia/cmap_algorithms.

The results represent the status of the CLUE platform when last accessed on June 4 to 5, 2019; the database is dynamic and changes slightly as new perturbagens are added.

## 3. Results

A portion of the heatmap output for an inquiry of gene expression signatures with similarities to PBZ is presented in Fig 2. Obviously, PBZ profiled against itself results in the highest score. It should be noted that the median tau (connectivity) score for the 9 cell results and the summary is reported in this output presentation. When searching this output in the cloud-based platform, the tau scores for the individual cell results can be accessed by placing the cursor over the cell result. This is an important feature of the software since it allows identification of the relative connectivity for a query perturbagen with each of the 9 tumor cell types, which may be valuable in recognizing specific mechanisms of action that may have therapeutic applications.

**Fig 2.**
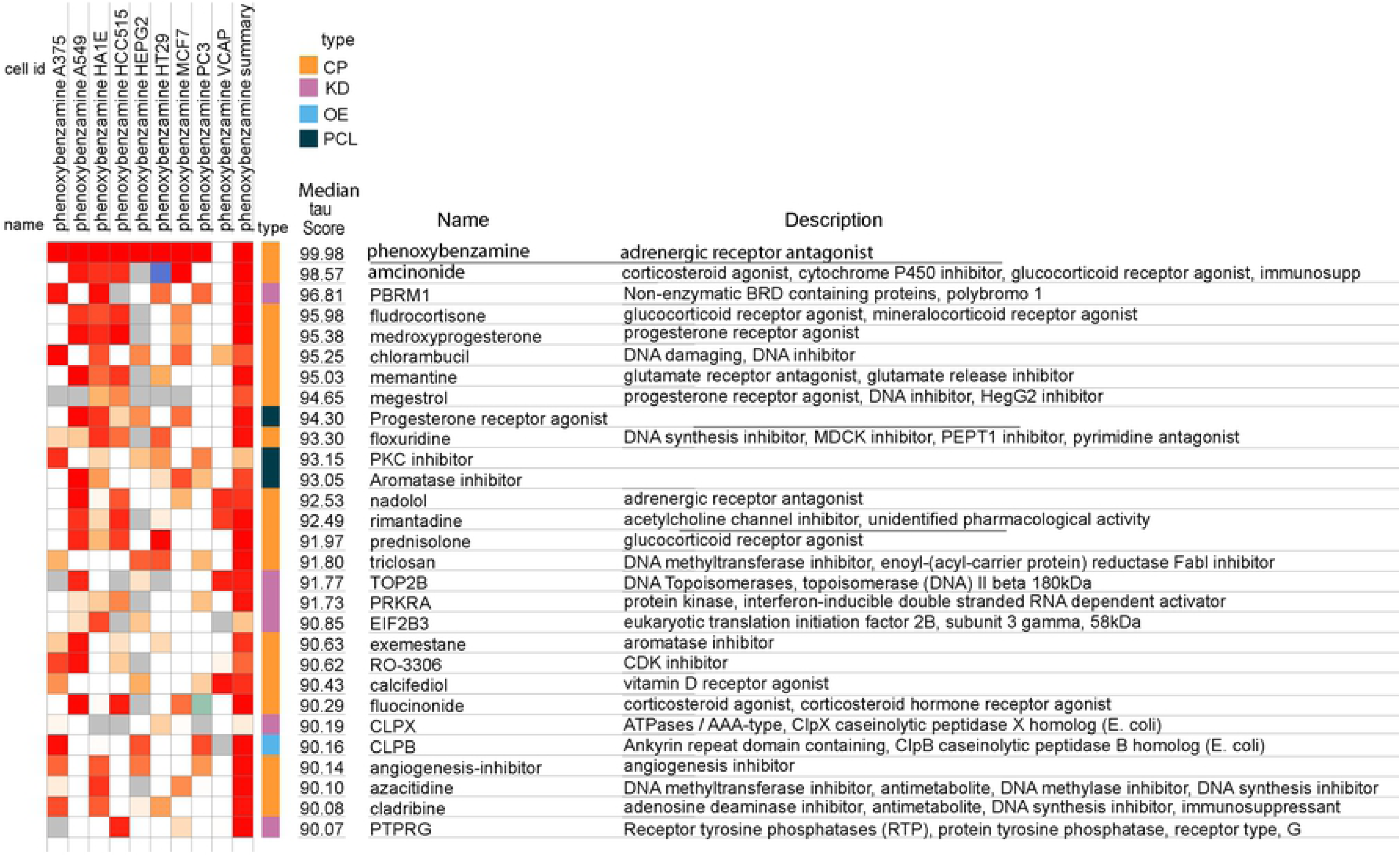
Heatmap of association strengths (median tau score) between gene expression signatures of phenoxybenzamine in 9 human malignant cell cultures and those of all entities in the Touchstone database. The cell identities (id) are listed in Table 1. The CMap classes (type) are as follows: Chemical compound/pharmacologic agent (CP); gene knock-down (KD); gene over-expression (OE); and perturbagen class (PCL).

For illustration purposes, the heat map screen shot in Fig 2 presents median tau scores that are 90 or greater. These results are generated from comparisons between PBZ and 8387 of the four CMap classes of perturbagens in the Touchstone database, i.e., CP, KD, OE, and PCL, as seen under the column labeled “type.” Even in this limited portion of the output, there are 4 glucocorticoid receptor agonists represented. As noted above, and illustrated further below, glucocorticoids ranked as the largest perturbagen class in relation to gene expression signatures that were similar to PBZ.

When the detailed list option in Touchstone was queried for gene expression signatures that had connectivity with PBZ it returned the gene set enrichment scores (2, 10) for groups of compounds represented in defined PCL sets (Table 2). A number of perturbagen classes of biological interest were identified with group scores above 90; these are considered by the program developers to be of strong interest in development of hypotheses for mechanisms of action and therapeutic application; see: https://clue.ie/connectipedia/connectivity_scores. There were no perturbagen classes that had group scores opposing the gene expression profiles of PBZ, i.e., there were none with scores that were −90 or lower. The perturbagen classes are organized initially on the basis of 3 or more perturbagens with the same mechanism of action. However, it is essential that there is strong connectivity of gene expression signatures among all members of a class; see: https://clue.io/connectopedia/pdf/pcls.

**Table 2.** CMap Class: PHARMACOLOGIC

| <u>Perturbagen Class (PCL)</u> | <u>Group Enrichment Score</u> |
| --- | --- |
| Progesterone receptor agonist x 12 | 99.50 |
| Glucocorticoid receptor agonist x 44 | 98.78 |
| Aromatase inhibitor x 3 | 97.98 |
| Glycogen synthase kinase inhibitor x 11 | 97.63 |
| HSP inhibitor x 7 | 96.76 |
| EGFR inhibitor x 14 | 96.33 |
| PPAR receptor agonist x 16 | 95.40 |
| DNA synthesis inhibitor x 3 | 95.02 |
| PKC inhibitor x 7 | 93.75 |
| JAK inhibitor x 5 | 92.59 |
| HIV protease inhibitor x 4 | 92.34 |
| Cannabinoid receptor agonist x 8 | 91.13 |

**HSP**, heat shock protein 90; **EGFR**, epidermal growth factor; **PPAR**, peroxisome proliferator-activated receptor; **PKC**, protein kinase C; **JAK**, Janus kinase; **HIV**, human immunodeficiency virus.

Each of the perturbagen classes in Table 2 was examined in the Touchstone detailed list for its members; these results are presented in Tables 3, 4, and 5. Only comparisons of PBZ with chemical compounds/drugs (the CP class) were searched in these queries; this represented 2429 agents. For these analyses, all agents with connectivity scores above 85 when compared with PBZ were included; for example, in the case of drugs with glucocorticoid agonist activity (Table 3) this included 30 of the 44 agents of that perturbagen class with this activity in the Touchstone database.

**Table 3.** Perturbagen Class with a Major Number of Gene Expression Signature Similarities to Phenoxybenzamine

| Score | ID | Name | Description | Targets |
| --- | --- | --- | --- | --- |
| ***Glucocorticoid Receptor Agonists*** |  |  |  |  |
| 99.93 | 0024 | Depomedrol | Glucocorticoid receptor agonist |  |
| 99.93 | 9173 | Halcinonide | Glucocorticoid receptor agonist | NR3C1 |
| 99.93 | 3476 | Clocortolone | Glucocorticoid receptor agonist | NR3C1, PLA2G1B |
| 99.93 | 8138 | Budesonide | Glucocorticoid receptor agonist | NR3C1, CYP3A5, CYP3A7 |
| 99.93 | 1468 | Prednisolone | Glucocorticoid receptor agonist | NR3C1, NR3C2, SERPINA6 |
| 99.93 | 5801 | Fludroxycortide | Glucocorticoid receptor agonist | NR3C1, SERPINA6 |
| 99.93 | 9782 | Fludrocortisone | Glucocorticoid receptor agonist | NR3C2, AR, NR3C1 |
| 99.93 | 8930 | Amcinonide | Glucocorticoid receptor agonist | NR3C1, ANXA1 |
| 99.89 | 0537 | Beclometasone | Glucocorticoid receptor agonist | NR3C1, CYP3A5, GPR97, SERPINA6 |
| 99.82 | 6322 | Fluocinonide | Glucocorticoid receptor agonist | NR3C1, SERPINA6, SMO |
| 99.72 | 0630 | Mometasone | Glucocorticoid receptor agonist | NR3C1 |
| 99.72 | 0871 | Triamcinolone | Glucocorticoid receptor agonist | NR3C1, CYP3A5, CYP3A7, SERPINA6 |
| 99.51 | 8384 | Beclometasone | Glucocorticoid receptor agonist | NR3C1, CYP3A5, GPR97, SERPINA6 |
| 99.44 | 3550 | Prednisolone | Glucocorticoid receptor agonist | NR3C1, NR3C2, SERPINA6 |
| 98.27 | 0379 | Fluticasone | Glucocorticoid receptor agonist | NR3C1, CYP3A5, CYP3A7, NR3C2, PGR, PLA2G4A |
| 98.27 | 3907 | Isoflupredone | Glucocorticoid receptor agonist | NR3C1 |
| 98.13 | 0232 | Westcort | Glucocorticoid receptor agonist |  |
| 97.74 | 6775 | Hydrocortisone | Glucocorticoid receptor agonist | ANXA1, NOS2, NR3C1, NR3C2 |
| 97.53 | 9987 | Flunisolide | Cytochrome P450 inhibitor | NR3C1 |
| 97.50 | 6607 | Flumetasone | Glucocorticoid receptor agonist | NR3C1, PLA2G1B |
| 97.38 | 5496 | Clobetasol | Glucocorticoid receptor agonist | NR3C1, PLA2G1B |
| 96.45 | 7533 | Rimexolone | Glucocorticoid receptor agonist | NR3C1, SERPINA6 |
| 95.26 | 3086 | Loteprednol | Glucocorticoid receptor agonist | NR3C1 |
| 93.94 | 3609 | Fluocinolone | Glucocorticoid receptor agonist | NR3C1, SERPINA6 |
| 89.54 | 7842 | Prednisolone | Glucocorticoid receptor agonist | NR3C1, NR3C2, SERPINA6 |
| 87.05 | 1694 | Alclometasone | Glucocorticoid receptor agonist | NR3C1, SERPINA6 |
| 86.97 | 4993 | Diflorasone | Corticosteroid agonist | NR3C1, PLA2G1B |
| 86.66 | 0903 | Betamethasone | Glucocorticoid receptor agonist | NR3C1 |
| 85.73 | 7903 | Prednicarbate | Phospholipase activator | NR3C1, PLA2G1B |
| 85.00 | 3711 | Hydrocortisone | Glucocorticoid receptor agonist | ANXA1, NOS2, NR3C1, NR3C2 |

**NR3C1**, glucocorticoid receptor; **NR3C2**, mineralocorticoid receptor; **PLA2G1B**, phospholipase A2 Group 1B; **CYP3A5**, **CYP3A7**, cytochrome P450 enzymes; **SERPINA6**, corticosteroid-binding globulin; **AR**, androgen receptor; **ANXA1**, annexin A1; **GPR97**, adhesion G protein-coupled receptor; **SMO**, smoothened, frizzled class receptor; **PGR**, progesterone receptor; **PLA2G4A**, cytosolic phospholipase A2; **NOS2**, nitric oxide synthase 2.

**Table 4.**
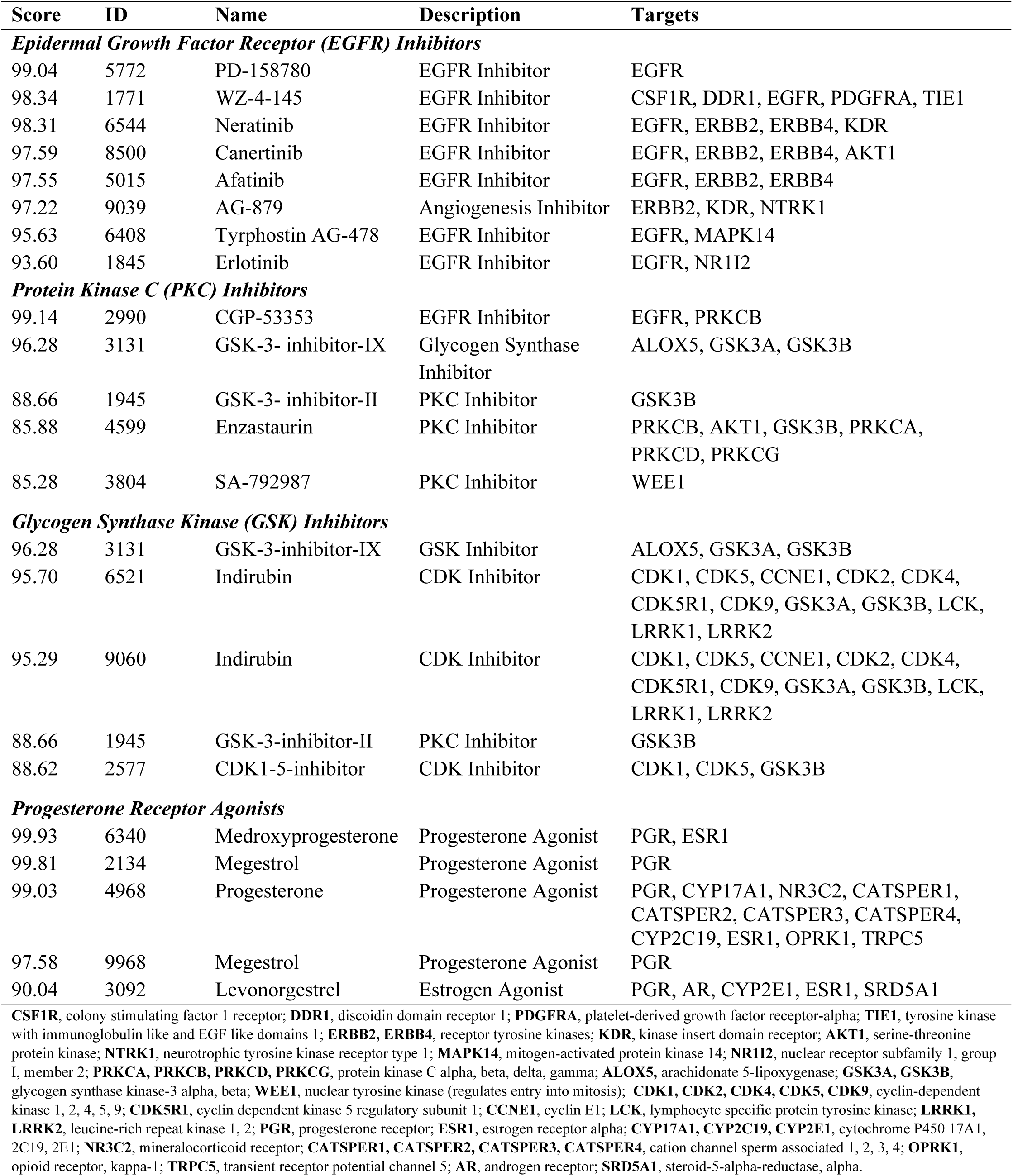
Perturbagen Classes with an Intermediate Number of Gene Expression Signature Similarities to Phenoxybenzamine

**Table 5.**
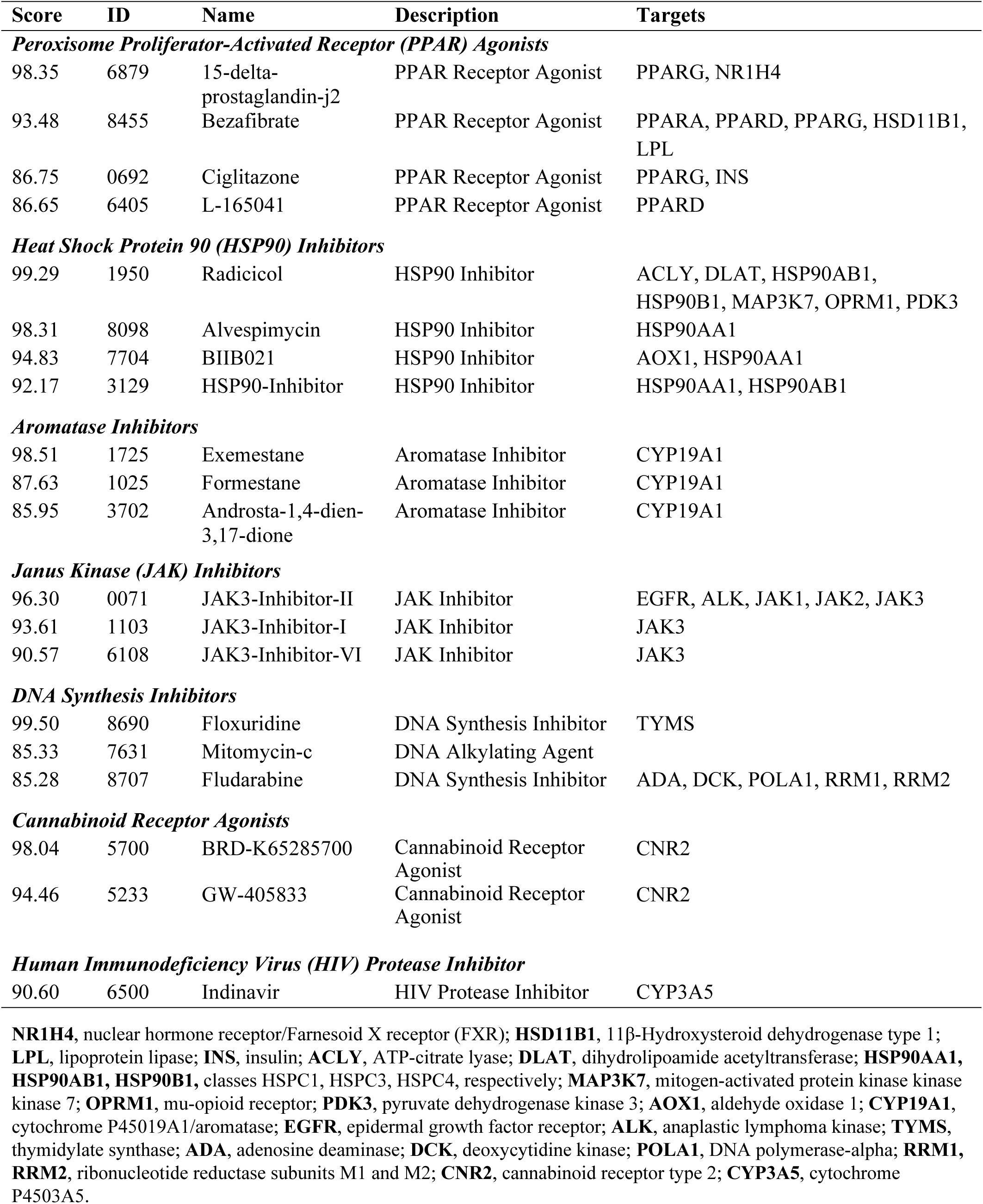
Perturbagen Classes with a Smaller Number of Gene Expression Signature Similarities to Phenoxybenzamine

HDAC inhibitors were not represented as a PCL set in the initial output (Table 2), but a search of this class in the detailed list database of Touchstone yielded the connectivities with PBZ for some of these agents (Table 6). Confirmation of connectivity of PBZ with HDAC inhibitors in the CLUE platform was consistent with our earlier report using the on-line version of CMap, and where PBZ was shown to directly inhibit several HDAC enzymes (1).

**Table 6.** Histone Deacetylase (HDAC) Inhibitors with Gene Expression Signature Similarities to Phenoxybenzamine

| Score | ID | Name | Description | Targets |
| --- | --- | --- | --- | --- |
| 96.77 | 2207 | Phenylbutyrate | HDAC inhibitor | CYP3A5, HDAC1 |
| 94.51 | 6356 | Rhamnetin | HDAC inhibitor | ALOX5, MAPK8 |
| 94.09 | 8671 | HNHA | HDAC inhibitor | HDAC1 |
| 91.58 | 9950 | NSC-3852 | HDAC inhibitor | HDAC1 |
| 91.00 | 7640 | JNJ-26854165 | HDAC inhibitor | HDACs1, 2, 3, 4, 5, 6, 7, 8, 9, 10, 11, MDM2 |
| 85.70 | 2742 | Trichostatin-A | HDAC inhibitor | HDACs1, 2, 3, 4, 5, 6, 7, 8, 9, 10 |

**CYP3A5**, cytochrome 4503A5; **ALOX5**, arachidonate 5-lipoxygenase; **MAPK8**, mitogen-activated protein kinase 8; **MDM2**, E3 ubiquitin-protein ligase *Mdm2*.

There are some prominent individual examples of gene expression connectivity between PBZ and other perturbagens in the database. One is a summarized connectivity score between PBZ and thalidomide of 98.52. This is of interest because both thalidomide and PBZ appear to be effective agents in the treatment of CRPS (11–15); also, thalidomide is an established treatment for myeloma and, in our earlier study, inhibition of growth in a myeloma cell line was one of the more potent effects of PBZ(1).

A second individual example was found in the connectivity score of 99.65 between PBZ and the glutamate receptor antagonist and glutamate release inhibitor, memantine. Memantine has been shown to have inhibitory effects on the growth of malignant gliomas (16) (17) and PBZ showed anti-proliferative effects on the glioblastoma malignant cell line, U-251, in our previous report (1). It can be seen that PBZ had a median tau score of 95.03 in comparison with memantine in the heatmap format (Fig.2), where it was compared with the entire Touchstone database of 8387 entities. As noted, the comparison between PBZ and memantine in the detailed list, with only 2429 chemicals and drugs, is the higher ranking value of 99.65. Thalidomide did not rank above 90 in the heatmap output.

## 4. Discussion

### 4.1 Glucocorticoid receptor agonists

Table 2 presents the summary of sets of agents by defined perturbagen classes that showed strong connectivity scores with PBZ. The most prominent class was glucocorticoid receptor agonists; this was completely unanticipated since PBZ shows no resemblance to a steroid chemical structure (Fig 1). There are a total of 44 drugs with glucocorticoid receptor agonist activity in the Touchstone database; the fact that so many individual glucocorticoids had connectivity scores of 85 or greater (Table 3) would indicate a strong similarity in their gene expression signatures to those of PBZ among the 9 cell types in the analyses. Of course, glucocorticoids have major systemic and topical therapeutic applications in inflammatory, autoimmune, and allergic syndromes(18); the current finding of their broadly similar gene expression signatures with PBZ is consistent with the anti-inflammatory/immunomodulatory mechanism of action for PBZ that has been proposed from some of our work and that of others in the neuropathic pain syndrome, CRPS (1, 14, 15, 19).

Also, it is possible that some of the apparent anti-proliferative effects of PBZ that were observed in malignant cell cultures (1) may have a basis in the glucocorticoid receptor agonist activity that is deduced from the CLUE platform. Glucocorticoids have established efficacy in the treatment of several hematopoietic malignancies with lymphatic lineage, including chronic lymphocytic leukemia, acute lymphoblastic leukemia, multiple myeloma, Hodgkin’s lymphoma and non-Hodgkin’s lymphoma (18) (20). Induction of apoptosis appears to be a primary mechanism in the treatment of these malignancies (21). In a connection with our previous report, PBZ showed some of its more potent anti-tumor effects against the human lymphoma cell line, SU-DHL-1(1)

The major drawback in the therapeutic use of glucocorticoids is the adverse effect profile for these drugs; major toxicities include immunosuppression, Cushing’s syndrome, osteoporosis and glucose intolerance. These effects are dose dependent and reductions in dosages lead to loss of therapeutic effectiveness. As noted, PBZ has no chemical relationship to a glucocorticoid steroid, and none of the classical steroid adverse effects have been observed in its clinical use since its approval by the United States Food and Drug Administration (FDA) in 1953. PBZ is classified chemically as a haloalkylamine (Fig 1). It is only FDA labeled for the treatment of hypertensive emergencies, most particularly for the control of surges of blood pressure from tumors of the adrenal medulla, termed pheochromocytomas. Its proprietary name in the United States is Dibenzyline, and there are generic preparations of the drug. The drug forms covalent bonds with α_1_- and α_2_ – adrenergic receptors resulting in a long-lasting non-competitive antagonism of these receptors. The drug has several non-FDA-labeled indications related to its relaxing effects on vascular smooth muscle in peripheral vascular diseases and the smooth muscle of the urogenital tract (http://www.ahfsdruginformation.com; Section 12:16.04.04) (22). Its vasodilating effect is responsible for most of its relatively minor dose-related side effects of hypotension, dizziness, nasal stuffiness and impotence; these effects appear to be self-limiting to some extent with continued intake (14, 15, 23).

The clinical use of PBZ for its smooth muscle relaxing effects on the urogenital tract is a source of information on the tolerability of the drug with long-term administration. This is found in particular with the earlier use of the drug for the treatment of the obstruction of urine flow in benign prostatic hypertrophy (BPH). Te (23) has summarized the results of 6 clinical trials with the use of PBZ for the chronic treatment of BPH; a total of 530 patients were represented in these trials and no reports of steroid hormone-like side effects were noted. The doses in these trials ranged from 10 to 40 mg per day. As noted above in Methods, the standard assay concentration for generation of gene expression profiles for perturbagens with the 9 tumor cell types was 10 µM; for the glucocorticoid data in Table 3, the 10 µM concentration was used throughout for all drugs, including PBZ. Although pharmacokinetic considerations about bioavailability for PBZ and specific glucocorticoids would need to be considered for any dosage comparisons between drugs, it is of interest in a general sense that a range of 10 to 40 mg per day includes chronic dosage levels for many clinically important glucocorticoids (18).

### 4.2 Progesterone receptor agonists

Progesterone receptor agonists are another class of steroid hormones that have applications for certain inflammatory syndromes and cancer therapy. Table 4 includes gene signature connectivities with PBZ for several important progesterone receptor agonists that are used in hormone therapy (24). The interaction of estrogen and progesterone receptors in malignant transitions, progression, and metastasis causes the application of therapeutic choices to be specific to the organs involved and the state of disease. Progesterone receptor agonists have a recognized place in the treatment of the hyperplasia and pain of endometriosis and in its possible progression to endometrial cancer, where progesterone agonists are used with some efficacy at various stages of the disease (http://www.ahfsdruginformation.com; Section 68:32) (22) (24) (20).

### 4.3 Epidermal Growth Factor Receptor Inhibitors

Eight gene expression signatures for inhibitors of epidermal growth factor receptor (EGFR) co-sorted strongly with PBZ (Table 4). EGFR inhibition has become a major area of investigation and clinical application for many cancers. Monoclonal antibodies directed against the EGFR receptor and tyrosine kinase inhibitors of these receptors are prominent in clinical treatment today, and other modalities to target and inhibit the receptor are under development (25). Three of the gene expression signatures for small molecule EGFR kinase inhibitors in Table 4, erlotinib, neratinib and afatinib, are part of the established formulary for neoplastic disease directed against EGFR receptors (26).

### 4.4 Protein Kinase C Inhibitors

Five gene signatures are identified for protein kinase C (PKC) inhibitor signatures that strongly co-sort with PBZ (Table 4). PKC is found in virtually all cells and tissues, with a large number of isoforms mediating a multitude of physiological and pathologic pathways. A number of clinical trials are at various stages with the use of inhibitors of specific isoforms for the prevention of the progression of malignancies and also for treatment of vascular changes in diabetes (27, 28). A recent extensive review covers the importance of PKC as a target for regulation in a number of vascular disorders (29). Among the several investigational PKC inhibitors listed in Table 4, the drug enzastaurin has recently been shown to have efficacy in improving survival in the treatment of diffuse large B-cell lymphoma (30) (31).

### 4.5 Glycogen Synthase Kinase (GSK) Inhibitors

There are 5 members in this perturbagen class that have gene signature enrichment scores above 85 in connectivity with PBZ (Table 4). The agent with the highest enrichment score relative to PBZ in this class, GSK-3-inhibitor-IX, is bromide substituted indirubin; indirubin assayed in 2 instances is the second and third highest matches with PBZ. GSK-3-inhibitor-IX shows primary activities as a GSK, PKC and lipoxygenase inhibitor; the primary activities of indirubin are as a cyclin dependent kinase (CDK), GSK and PKC inhibitor. The protein targets for these inhibitors show additional overlap.

CDK exists in a number of isoforms. CDK1 expression is elevated in a number of human cancers, such as Hodgkin’s lymphomas, cancer of the prostate, gastric lymphoma, lymphoblastic leukemia, gastric lymphoma and ovarian cancer. Inhibition of CDK1 shows suppression of epithelial ovarian cancer growth and the effectiveness is enhanced by co-treatment with other anti-cancer agents, such as cisplatin (32). There is an extensive literature on the anti-proliferative effects of indirubin and its derivatives; it is a natural product that has been used in traditional Chinese medicine (33). One of its earliest clinical applications was in the promising treatment of chronic myelocytic leukemia (34). There is continued research on its potential in other malignancies, psoriasis, pulmonary arterial hypertension, and restenosis (33).

### 4.6 Peroxisome Proliferator-Activated Receptor (PPAR) Agonists

Four gene expression signatures for PPAR agonists co-sorted strongly with gene expression for PBZ (Table 5). PPAR agonists are an active area of research and clinical development because activators of these receptors result in an improvement of insulin sensitivity in the treatment of type 2 diabetes (35). Two of the perturbagens that are identified with gene expression signatures similar to PBZ, bezafibrate and ciglitazone, have known clinical importance. Bezafibrate has been studied in a number of clinical trials (36) (37); it is considered to possibly be superior to other fibrates in that it has broad agonist activity on all PPAR isoforms, alpha, gamma and delta. Selective PPAR gamma agonists, such as the thiazolidinediones that are used today in the treatment of Type 2 diabetes mellitus, have the adverse effect of causing water retention and weight gain, an effect that is not prominent with bezafibrate (36).

The second PPAR agonist of recognized clinical importance that matched with PBZ gene expression is ciglitazone. Ciglitazone is the chemical prototype of all thiazolidinediones although it was not considered clinical efficacious enough to be marketed (38). In addition to the lead that this drug provided to the development of clinically important drugs for the treatment of Type 2 diabetes, ciglitazone was found experimentally to have in-vitro and in-vivo anti-tumor activity in melanoma (39) and glioblastoma (40). Research continues on the potential of thiazolidinediones for their anti-bacterial activity and anti-proliferative effects in neoplastic diseases (41).

### 4.7 Aromatase Inhibitors

Table 5 presents data for the connectivity scores of the gene expression signatures for PBZ with those for 3 aromatase inhibitors. All 3 agents are steroidal inhibitors of aromatase, which is CYP19A1. This enzyme is responsible for the conversion of androgens to estrogens, and inhibition results in a lowering of estrogens levels (20). The first two agents, exemestane and formestane have proven clinical efficacy in the treatment of hormone-receptor positive (HR+) breast cancer. Formestane is no longer used clinically because it required injection, while exemestane is orally active. These drugs bind irreversibly to aromatase resulting in a “suicidal” type inhibition (42). Exemestane represents the 3^rd^ generation class of aromatase inhibitors and has wide application. It is used for initial early-stage treatment of breast cancer in HR+ postmenopausal women and for more advanced and metastatic cancer. It is sometimes used in combination treatment with a CDK4/6 inhibitor. Third generation aromatase inhibitors are also used as breast cancer prevention therapy in post-menopausal women (20).

### 4.8 Janus Associated Kinase (JAK) Inhibitors

Three gene signatures for Janus associated kinase inhibitors were identified for similarity to PBZ signatures (Table 5). Janus kinases are downstream signaling mediators for a number of cytokines that result in inflammatory processes associated with, for example, inflammatory bowel disease, rheumatoid arthritis, psoriasis, etc. They show significant promise for the treatment of inflammatory and immune pathologies (43).

The three JAK inhibitors in Table 5 that co-sorted with PBZ are experimental agents; they are chemically identified as follows:

Broad Institute I.D. 0071 JAK3-Inhibitor-II 4-[[6,7-bis(hydroxymethyl)-4-quinazolinyl] amino]-2-bromophenol.

Broad Institute I.D. 1103 JAK3-Inhibitor-I 4-[[6,7-bis(hydroxymethyl)-4-quinazolinyl]amino]phenol.

Broad Institute I.D. 6108 JAK3-Inhibitor-VI 5-(3-pyridinyl)-3-(1H-pyrrol-2-ylmethylidene)-1H-indol-2-one.

A recent report summarizes the 2 approved drugs in this class, tofacitinib and decernotinib, and the extensive clinical testing currently in progress of 8 additional JAK inhibitors. They are being developed for the treatment of ulcerative colitis, Crohn’s disease, atopic dermatitis, alopecia areata, diffuse scleroderma, vitiligo, infectious uveitis, cutaneous lupus erythematosus and hemophagocytic syndrome (44).

### 4.9 DNA Synthesis Inhibitors

Three DNA synthesis inhibitors, floxuridine, mitomycin-C and fludarabine showed connectivity with PBZ with enrichment scores greater than 85 (Table 5). All are currently clinically important drugs in the treatment of neoplastic diseases (45). The fact that PBZ had connectivity with these drugs prompted inspection of the PCL sets that were identified when floxuridine was entered as the Index compound in the Touchstone database. Six of the 12 PCLs identified for PBZ (Table 2) were also identified for floxuridine; they were, in rank order of enrichment score: DNA synthesis inhibitor; Progesterone receptor agonist; PPAR receptor agonist; Cannabinoid receptor agonist; Glycogen synthase kinase inhibitor; and JAK inhibitor. It would appear that there is the possibility of a number of similarities between the gene expression signatures of PBZ and DNA synthesis inhibitors that could account for the connectivity scores for these drug and common anti-proliferative effects that they may share. As with so many cancer chemotherapeutic drugs, the DNA synthesis inhibitors have serious adverse effects. It is important to note a finding that we have not previously reported. In the effects of PBZ on human cancer cell growth assays that were conducted for us by the National Cancer Institute (NCI60), and reported in our previous study(1), PBZ showed no lethality in any of the cell cultures at their standard assay concentration of 10 µM.

### 4.10 Heat Shock Protein 90 Inhibitors

There are 7 gene signatures for Heat Shock Protein (HSP90) inhibitors that rank with that for PBZ (Table 5). HSP90 inhibitors represent a current intense area of clinical investigation as adjunctive agents for the treatment of multiple cancers. Radicicol, the highest scoring agent in relation to PBZ (Table 5), has been reported to induce apoptosis in malignant human epithelial ovarian cells (46) and a phase I study with alvespimycin in patients with acute myeloid leukemia gave preliminary evidence of efficacy (47). There is also a more recent trial with alvespimycin for the treatment of relapsed chronic lymphocytic leukemia and small lymphocytic lymphoma (48).

Wang et al. has compiled a review of 15 published phase II clinical trials with HSP90 inhibitors (49). They represent treatments in 10 different types of cancer where the majority of the patients had relapsed after prior treatment with other agents. One of the agents tested in refractory cases of gastrointestinal stromal cancer was BIIB021, the third agent with a gene expression signature similar to PBZ (Table 5).

### 4.11 Cannabinoid Receptor Agonists

Table 5 lists 2 experimental “selective” Type 2 (CB2) cannabinoid receptor agonists that showed strong gene expression connectivities with PBZ. The first compound, with a Broad Institute identification of BRD-K65285700, is N-(piperidin-1-yl)-1-(2,4-dichlorophenyl)-1,4-dihydro-6-methylindeno(1,2-c)pyrazole-3-carboxamide. There is interest in the development of drugs with selectivity for the CB2 receptor in an attempt to optimize the potential therapeutic effects of the cannabinoids in regard to pain alleviation, anti-inflammatory effects and anti-cancer effects and avoid the undesirable central nervous system psychoactive effects related to agonist activity on CB1 receptors (50). However, the role of CB2 receptors in the brain in promoting anti-nociceptive and behavioral effects has advanced as well (51). Research with the second agent, GW-405833, has progressed considerably with evidence for treatment of inflammatory and neuropathic pain (52). Of interest, amelioration of neuropathic pain appears to also involve a non-competitive antagonism of CB1 receptors by GW-405833 (53). In this connection, the fact that cannabinoids show gene expression signatures similar to PBZ is of interest in regard to the observations of apparent efficacy of PBZ in the treatment of neuropathic pain syndromes (15).

There is considerable interest in the potential for cannabinoids as anti-cancer drugs (54) (55). These studies are essentially limited to in-vitro models and rodent experiments. Daris et al. have also reviewed the observation that some of the plant-derived cannabinoids have the potential for tumor proliferation at nanomolar concentrations in certain cell lines while showing a biphasic effect of tumor suppression at micromolar concentrations. This is an obvious consideration in terms of the choice of agent and the clinical context in its use.

Endogenous, plant-derived and synthetic cannabinoids are also being evaluated for efficacy in several neuro-inflammatory and degenerative diseases. There is both preclinical and clinical trial data available. Baul et al. (56) have focused primarily on research in Parkinson’s disease; Chiurchiu et al.,(57) have reviewed hypotheses and studies of cannabinoids in relation to multiple sclerosis, Alzheimer’s disease, Parkinson’s disease, Huntington’s disease, and amyotrophic lateral sclerosis.

### 4.12 HIV Protease Inhibitors

PBZ showed significant gene expression connectivity with one HIV protease inhibitor, indinavir (Table 5). As with other HIV protease inhibitors, indinavir has a peptide similar structure and is a competitive inhibitor of aspartyl protease. It is currently in clinical use but is used less frequently today since it is not superior to other inhibitors of HIV protease and has more concerns related to renal toxicity (58).

In an attempt to gain some insight into possible reasons for the overlap between gene expression signatures for PBZ and indinavir, indinavir was queried as the index perturbagen in Touchstone. The PCL sets that were returned as important for indinavir relative to the entire database were not similar to any of those for PBZ (Table 2). However, there were two PCL sets for indinavir, “Norepinephrine reuptake inhibitor,” and “Tricyclic antidepressant” that had high enrichment scores. Inhibition of norepinephrine reuptake in the central nervous system is believed to be involved in the therapeutic effects of the tricyclic antidepressants. This is of interest because PBZ is known to be a potent inhibitor of both neuronal and extra-neuronal uptake of norepinephrine (59).

### 4.13 Histone Deacetylase Inhibitors

The HDAC inhibitor with the highest connectivity score to PBZ was phenylbutyrate (Table 6). Phenylbutyrate is designated as an orphan drug by the FDA with applications in the treatment of promyelocytic leukemia, primary or recurrent malignant glioma, amyotrophic lateral sclerosis, and spinal muscular atrophy. The first two indications are approved by a major United States medical insurance company; “Aetna considers sodium phenylbutyrate medically necessary for the treatment of acute promyelocytic leukemia and malignant glioma.” (http://www.aetna.com/cpb/medical/data/200_299/0240.html) This link provides an extensive literature review of the research on other potential therapeutic indications for phenylbutyrate.

The next 4 HDAC inhibitors listed in Table 6 have various stages of experimental investigation. Rhamnetin has been investigated for its anti-inflammatory effects; at a concentration of 1 µM it caused significant inhibition of pro-inflammatory cytokines from macrophages (60).

N-hydroxy-7(2-naphthylthio) heptanomide (HNHA) was found to be more potent than trichostatin-A (TSA) and vorinostat (SAHA) in inhibition of proliferation of anaplastic and papillary human thyroid cell cultures; HNHA also increased histone acetylation in both of these cells types and reduced tumor volume in athymic nude mice and prolonged survival to a greater extent than either TSA or SAHA (although all three HDAC inhibitors showed efficacy in these regards) (61).

Martirosyan et al. found that NSC-3852, a quinoline compound, inhibited HDAC activity in-vitro, induced differentiation, and caused DNA damage and apoptosis in MCF-7 human breast cancer cells (62). This same laboratory found that the cell differentiating effects and apoptosis were due to the generation of reactive oxygen species (63).

JNJ-26854165 is also known by its proprietary name, Serdemetan. It was extensively investigated preclinically for efficacy against a number of tumors and in a Phase I tolerability study; it did not proceed to a Phase II study (64). However, preclinical studies have continued for efficacy in several lymphomas (65), and most recently specifically for Burkitt’s lymphoma associated with the Epstein-Barr virus (66).

Trichostatin-A (Table 6) is a classic example of an experimental HDAC inhibitor with broad activity against a number of HDAC enzymes; its summarized score of 85.70 was consistent with the threshold for inclusion that we set. However, two FDA-approved HDAC inhibitors, belinostat, for treatment of peripheral T-cell lymphoma, and SAHA (Vorinostat), another classic HDAC inhibitor and approved for cutaneous T-cell lymphoma, had scores of 76.10 and 67.83, respectively (67, 68).

The overlap between the gene expression signatures of PBZ in the 9 malignant cell panel of the CLUE CMap platform and a number of HDAC inhibitors is of particular interest in relation to the original report from the earlier CMap platform that predicted HDAC inhibitory activity for PBZ (1). Follow up assays demonstrated that PBZ had essentially comparable inhibitory potency to trichostatin-A on HDAC5 and 9 and greater inhibitory potency than SAHA on both HDAC5 and 9. The selectivity of PBZ for inhibition of these HDACs is of interest in relation to the fact that the high expression of HDAC5 and 9 in pediatric medulloblastoma tumors was found to be prognostic for poor survival (69). These tumors are the most common pediatric brain tumors. HDAC5 has also been shown to be a strong promotor of tumor migration and invasion in hepatocellular carcinoma (70) and breast cancer (71). And, particular elevation of HDAC9 has been reported in cervical cancer, childhood acute lymphoblastic leukemia, Philadelphia-negative chronic myeloproliferative neoplasms and Waldenstrom’s macroglobulinemia (72), all raising the possibility for therapeutic trials of PBZ with its minimal side effect profile (14, 15).

### 4.14 Evidence of Anti-proliferative Activity of PBZ from Other Studies

Studies of PBZ anti-proliferative activity by other investigators, which were discussed in detail in an earlier publication (1), can be summarized as follows:

-- From screening of 1290 small molecules in human neuroblastoma cell lines, PBZ was one of four compounds that showed dose-dependent inhibition of growth, invasion and migration (73).

-- Cobret et al. found that PBZ showed the most specific and highest extent of complex formation with LINGO-1 from a screen of 1263 small molecules (74). This ligand appeared to promote the capacity of LINGO-1 to inhibit signaling through EGFR activation in neuronal elements, which may have relevance to a possible role in the inhibition of aggressive cancers.

-- In connection with the study by Cobret et al. (74), Lin et al. (75)showed that PBZ inhibited the growth, invasion and migration in U251 and U87MG glioma cell cultures and decreased tumor volume expansion *in-vivo* in mice with implanted U87MG cells that received direct injection of 20 nM PBZ into the tumors. These effects were accompanied by a marked increase in LINGO-1 expression following treatment with PBZ, which may have a relationship with the above observations.

-- Lee et al. (76) found that PBZ was able to substantially decrease the vascular proliferative activity that results in development of the pulmonary arterial hypertension associated with injection of monocrotaline, a pulmonary toxin. Compared with controls, medial thickening was decreased by 80% after 4 weeks of daily PBZ treatment that followed the monocrotaline challenge. Also, Kim et al. (77) found that specific HDACIIa inhibition significantly reversed established pulmonary arterial hypertension produced by both monocrotaline or hypoxic interventions.

Two recent studies have further advanced the knowledge of the anti-proliferative potential of PBZ. Zador et al. carried out a drug repurposing investigation utilizing Broad Institute CMap in search of therapeutic candidates for the treatment of atypical meningiomas (78). Compared with benign meningiomas, atypical tumors are characterized by a high risk of recurrence after surgery and radiation and a poor prognosis. Their analysis identified 3 drugs, verteporfin, emetine and PBZ, with the highest gene expression enrichment scores suggestive of opposition (potentially inhibitory/therapeutic) to progression of atypical meningiomas. PBZ was only 5% less strongly inhibitory than verteporfin; emetine placed second among the three (78).

Potential therapeutic applications for PBZ were also identified in an extraordinary recent drug repurposing study by Lin et al. that screened 640 FDA-approved drugs for their efficacies and potencies as macropinocytosis inhibitors (79). Macropinocytosis is distinct from receptor-mediated endocytosis and phagocytosis in that it is not mediated physically by interaction with the plasma membrane. And, as the term implies, it represents a much larger volume of engulfing of plasma content than either receptor-mediated endocytosis and phagocytosis; it has been described as a cell “drinking” process. Macropinocytosis has been implicated in a number of pathological syndromes. For example, cells involved in malignant proliferation engulf proteins en masse by macropinocytosis to support metabolic growth demands (80). Also, macropinocytosis has been implicated in the uptake of amyloid precursor protein in the development and progression of Alzheimer’s disease (81). In their drug screen of potential therapeutic inhibitors of macropinocytosis, Lin et al. limited their final candidates to drugs that did not interfere with physiologically important processes of receptor-initiated endocytosis or phagocytosis or toxicity considerations. This led finally to two candidates, the tricyclic antidepressant imipramine and PBZ (79). Although PBZ had a somewhat greater potency than imipramine in inhibiting the phorbol ester stimulation of dextran uptake by RAW 264.7 macrophages (IC_50_ = 43.8 nM and 130.9 nm, respectively), both agents would appear to be important for further development as inhibitors of pathological processes facilitated by micropinocytosis.

## 5. Conclusion

The present study is an extension of a previous report that suggested that the gene expression signatures that PBZ produced in two malignant cell lines were consistent with the possibility that it would have HDAC inhibitory activity. Investigation of PBZ in the greatly expanded Broad Institute CLUE platform revealed a number of previously unrecognized similarities between PBZ gene expression signatures and those of agents that would be consistent with potential antineoplastic, anti-inflammatory and immunomodulatory activity for PBZ. CMap was designed as a hypothesis-generating platform to help to identify possible relationships amongst gene-expression signatures in diseases and those produced by therapeutic interventions; however, the significance of such relationships must be confirmed by further biological experimentation. The fact that PBZ is an FDA-approved drug with a long clinical use that demonstrates its minimal adverse effect profile should facilitate further investigations into the possible therapeutic potential of the drug.

## Acknowledgement

The author is grateful to Ms. Jennifer Brown, Department of Pharmacology, New York Medical College, for editorial assistance in the preparation of the manuscript.

